# A redundant encoding algorithm for artificial sensory information speeds learning and improves multisensory-guided navigation

**DOI:** 10.64898/2026.09.17.752429

**Authors:** Samuel J. Senneka, Maria C. Dadarlat

## Abstract

Intracortical microstimulation (ICMS) directly modulates cortical activity, providing an artificial sensory stream to guide accurate control of prosthetic limbs. Yet, only a fraction of reported sensations evoked by ICMS are proprioceptive (i.e., describing the position and movement of the body). Taking a learning-based approach to encoding an artificial proprioceptive signal bypasses this limitation. For example, animals trained with multi-channel ICMS in association with natural vision learn to both decode the ICMS signal to localize an invisible target and learn to integrate artificial sensation with natural sensation. The limitation of a learning-based approach, however, is that it requires training. We hypothesized that changing the algorithm used to encode artificial sensory information could both reduce learning time and improve plateau performance on ICMS-guided navigation. To test this idea, we trained eight mice on a sensory-guided navigation task: mice were required to locate a target in a training cage, the position of which was encoded by a red circle (vision), by multi-channel ICMS (ICMS trials), or by both vision and ICMS (multi-modal). ICMS was encoded using one of two algorithms: sparse (fewer simultaneously stimulating electrodes) or redundant (more simultaneously stimulating electrodes). We found that redundant encoding sped learning of the ICMS signal relative to sparse encoding. Further, mice guided by redundant ICMS ran faster and completed trials in less time than when guided by sparse ICMS. Having redundant encoding also facilitated multisensory integration of ICMS with natural vision, improving success rates, path efficiency, movement speed, and movement time. We conclude that optimized ICMS encoding algorithms could overcome the current limitations of learning-based sensory encoding, facilitating learning and integration of artificial sensory information into existing sensorimotor neural circuits, paving the way to restoring both the sensory and motor streams of information flow in damaged sensorimotor systems.

## INTRODUCTION

Improving the quality of life for more than five million Americans who suffer from paralysis requires restoring both the ascending (sensory) and descending (motor) pathways of information flow [1, 2]. In cases where repairing severed connections is not possible (e.g., [1, 3–5]), sensorimotor function can be restored by establishing an artificial link between the brain and the periphery via neural prostheses (e.g., [6–9]). The essential elements of artificial sensory feedback from a prosthetic limb include tactile [10, 11] and proprioceptive sensations [12]; however, the development of artificial proprioceptive feedback for brain-machine interfaces (BMIs) has lagged, despite its clear benefit to BMI control [13]. Artificial proprioception would allow movements in the absence of vision and would be integrated with visual information when both are present to improve movement accuracy and precision [14–17].

An artificial sense of proprioception can be encoded using intracortical microstimulation (ICMS)

— passing small electrical currents through cortically-implanted electrodes to mediate artificial sensations [12, 18–23]. Multi-channel ICMS permits directly “writing-in” of information to the brain, guiding goal-directed movements [24–29]. While ICMS can evoke naturalistic sensations [11, 21, 23, 30–34], evoked proprioceptive percepts constitute a minority of evoked sensations and have low resolution for describing desired movement parameters [12]. We posit that, even with next-generation microelectrode arrays, encoding reliable and precise artificial proprioceptive signals will require learning on the part of the patient [35, 36].

The neural circuits underlying natural sensory processing remain plastic into adulthood [37– 39], including retaining the ability to develop multisensory responses to spatiotemporally-congruent sensory streams [40–42]. Multisensory integration — the process by which multiple sensory representations of a single variable, event, or object, are combined into a unified, and more precise, sensory estimate [15] — improves natural sensory-guided behavior [14, 16]. A similar facilitation occurs in *artificial* sensorimotor systems, including learned associations: mice trained on multi-sensory information (vision and ICMS of primary somatosensory cortex) rapidly learn to integrate natural and artificial sensation, significantly improving task performance [43].

A prevailing challenge in developing learning-based algorithms for encoding artificial sensation is identifying the key elements of an ICMS encoding algorithm that convey precise and reliable artificial sensory signals. There are clearly some constraints on learning artificial sensation, including anatomical: both patterned optogenetic and electrical stimulation are most easily learned when similar stimuli activate somatotopically-proximal brain regions [27, 44–47], ideally delivered to mid-to-deep layers of cortex [48]. Having more discrete encoding channels also helps encode finer information [26]. In addition to spatial considerations, higher stimulation frequencies are more readily detectable [49]. Finally, there is a cognitive component to learning: pre-training on a behavioral task guided by natural sensation speeds up learning of the same task guided by ICMS [26].

In this study, we test a theory, inspired by natural neural encoding of sensory information, that in many contexts, selective (sparse) encoding of behavioral variables is easier to learn and interpret than a distributed (redundant) code [50–52]. In the context of ICMS, a sparse code refers to isolating the encoded signal to one or two electrodes, while a redundant code refers to stimulating across multiple electrodes simultaneously. In both cases, the total number of possible stimulating channels remains the same [26].

We previously showed that mice receiving sparse, patterned multi-channel ICMS of mouse primary somatosensory cortex can encode the location of an invisible target [43]. The ICMS encoding algorithm was inspired by tuned responses of single neurons in primary somatosensory cortex (S1) to limb movements and tactile motion [24, 53–55]. We implemented redundancy by changing the width of a single electrode’s response distribution to the encoded variables (Figure 1A,B,D). Increasing tuning width has two effects on the signal: 1) the change in frequency per unit change in the encoded sensory variable decreases for each electrode, and 2) the number of electrodes stimulating to encode a value of the sensory variable increases (Figure **??**). While the former change reduces signal quality [52], the latter might enhance it.

**Figure 1:**
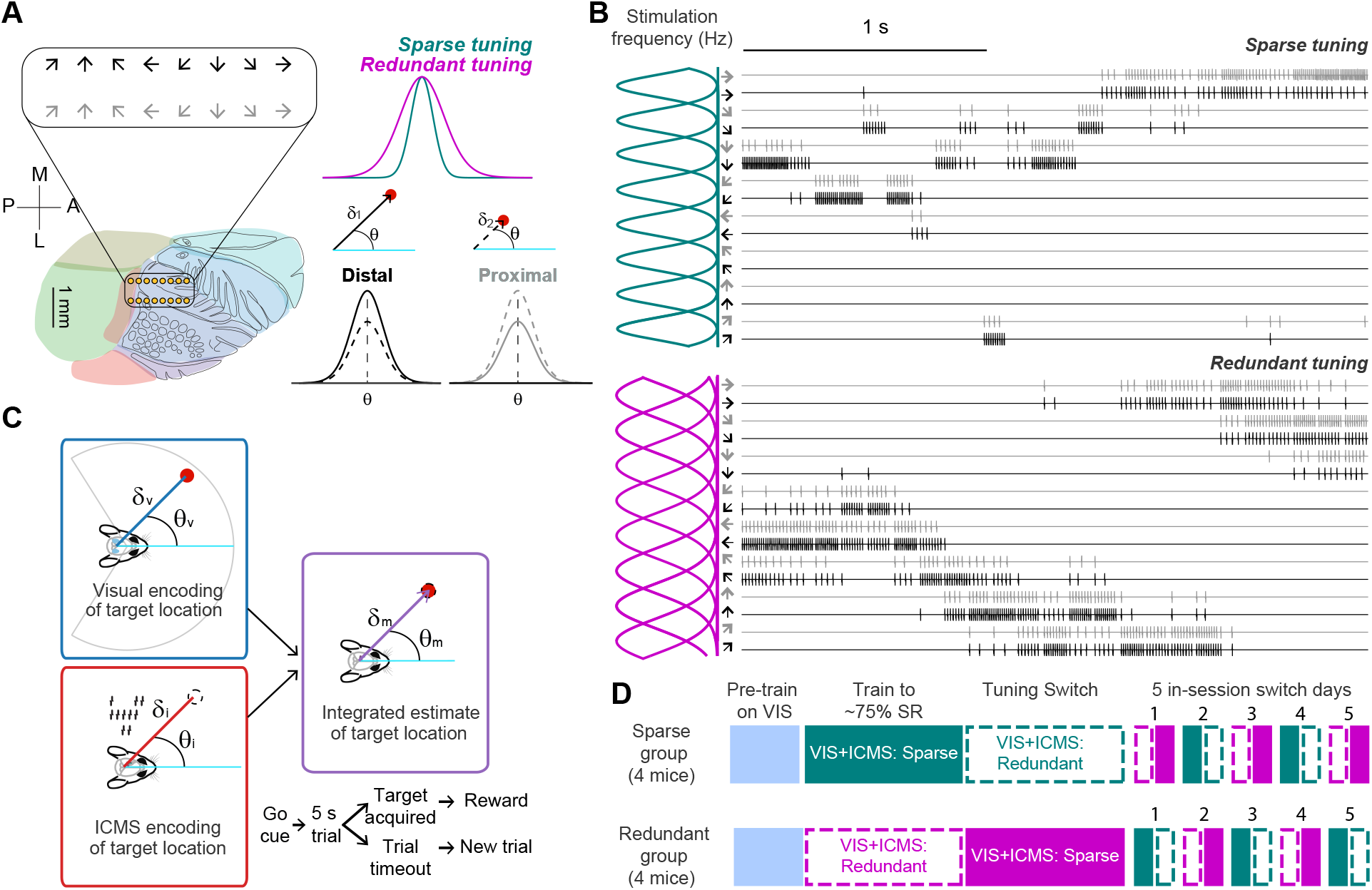
Experimental design. **A.** 16-channel microwire arrays arranged in an 8×2 grid were implanted over the forelimb/hindlimb region of mouse primary somatosensory cortex in the right hemisphere at a depth of 500 *µ*m (approximately layer V). The difference in tuning bandwidth for each electrode between sparse and redundant encoding algorithms (upper right). Rows were assigned to be distal or proximal encoders, with distal electrodes decreasing in maximum stimulation frequency as distance from the target decreases and proximal increasing as target distance decreases (lower right). **B**. Example stimulation traces for sparse (*top*) and redundant (*bottom*) encoding schemes. Curves along the y-axis show tuning for pairs of electrodes as well as overlap between electrode pairs for both encoding schemes. **C**. Schematic for integration and task structure. *Integration* Mice process visual and ICMS input to develop unimodal egocentric estimates of target location that are then combined into a multimodal estimate. *Task* Trial starts are indicated with an auditory go-cue, after which, mice have 5 seconds to navigate to the target region. Successful trials are rewarded with a small juice droplet, and failed trials result in a timeout period before a new trial starts. **D**. Training structure and timeline. All mice were pre-trained on the task with visual feedback. Mice were then assigned to sparse or redundant tuning groups and trained to proficiency on the task with ICMS feedback before switching to the other tuning width. Finally, mice were given 5 sessions in which tuning width was switched in the middle of each session, and the starting tuning width alternated by session.

Contrary to our initial hypothesis, we found that the redundant encoding scheme significantly sped learning of the ICMS sensory signal relative to sparse encoding. Within a behavioral trial, mice reached targets faster guided by redundant ICMS compared to sparse ICMS. The benefits of the ICMS encoding algorithm extended beyond decoding the ICMS signal. During multimodal trials, mice performed better with redundant encoding than with sparse encoding on all assessed metrics (success rate, time-to-target, average speed, angular dispersion), suggesting that redundant encoding facilitates the integration of artificial sensory information with natural sensation. Finally, mice were able to rapidly switch between sparse and redundant encoding schemes with little disruption to behavioral performance, showing remarkable flexibility in decoding ICMS.

We note that these experiments only probe behavioral performance, without considering the perceptual quality of evoked sensations. It is possible that modulating redundancy modulates the perceptual quality of the stimulation signal [56]. Even so, these results highlight that encoding algorithms can, and should, be optimized to integrate into the existing neural circuits that govern a broad suite of sensorimotor activities, thus paving the way for rapid, robust, and naturalistic sensory neural prostheses.

## METHODS

All procedures were approved by the Institutional Animal Care and Use Committee at Purdue University. Surgical methods and the behavioral task used for this experiment are described in detail in the Supplemental Extended Methods. Eight C57BL/6J mice aged three to four months were used in these experiments (four male, four female).

### Surgical Procedures

Each mouse was implanted with a 16-channel microwire array over the forelimb/hindlimb region of primary somatosensory cortex (Figure 1A), identified via stereotaxic coordinates [57]. Ground and reference wires were implanted in the contralateral hemisphere, just underneath the skull. Mice were given five to seven days to recover before training began.

### Behavioral Training and Testing

Mice were trained to navigate to targets located upon the floor of a custom behavioral training arena, guided by vision and/or ICMS signals that encoded the relative distance and direction between the mouse’s heading and the center of a target (Figure 1C). The target could appear in one of nine locations (Figure **??**), and mice were rewarded for moving their head over the target, for which they would receive a liquid reward from a reward port on the side of the chamber. Mouse head position and heading direction were tracked in real-time during the task using a camera positioned above the cage. The video was analyzed in real time using a DeepLabCut pose estimation model and the DeepLabCut-Live GUI [58]. Mice were cued to the start of the trial via an auditory tone, after which they would have five seconds to locate the target (Figure 1C). Failure to locate the target within the trial window would cause a timeout period.

A single behavioral session consisted of 150 training trials. Initially, trials were split as: 80% multimodal, 7.5% ICMS, 7.5% dim visual, and 2.5% for bright visual and sham each. No sensory information was provided about target location in sham trials, which provide a baseline performance level achievable by chance. Once mice reached proficiency on ICMS trials for a single day (defined as success rate *≥*75%), multimodal trials were decreased to 60%, visual were increased to 20% (15% dim visual target, 5% bright visual target), and ICMS was increased to 15%. The remaining 5% of trials were sham trials.

To compare learning of each encoding algorithm, mice were first randomly assigned to either sparse or redundant encoding. Of the eight mice included in the study, half were first trained on sparse encoding (N=4: 2 M, 2 F) and half were first trained on redundant encoding (N=4: 2 M, 2 F). The encoding algorithms used for each are described below in Equation 1. Three mice were trained in a first cohort; five mice in a second cohort.

Once mice were, on average, proficient on multimodal and unimodal ICMS trials across five days, they were switched to the alternative encoding algorithm (sparse to redundant, redundant to sparse) and trained for an additional five to ten sessions. We analyzed performance between the five sessions directly preceding and following the switch.

A final experiment was to test flexibility in decoding the two algorithms. To do so, mice were tested for five sessions in which the encoding algorithm was switched in the middle of the session. Mice began with the encoding algorithm used at the end of the previous session; after ∼75 trials, the encoding algorithm was switched. This experimental design controls for the order of presentation of the tuning widths within sessions.

### Intracortical Microstimulation (ICMS)

ICMS encoded the distance and direction between a mouse’s heading and the center of the target (Figure 1C) via patterned stimulation across all sixteen electrodes on the array (Figure 1A,B). Stimulation across each electrode consisted of a time series of cathode-leading biphasic pulses. Stimulation amplitude was kept fixed throughout training (10-20 *µ*A). Stimulation frequency was updated at 10 Hz (calculated via Eqn. 1) to reflect the changing position and heading of the mouse relative to the target.

Stimulation frequency for each electrode was modeled as a von-Mises function. First, each electrode was assigned a “preferred” direction that would represent a particular angle between the mouse’s forward heading and the direction of the target. Second, electrodes were assigned to be “distal” or “proximal”, which linearly decreased or increased maximum stimulation frequency as a function of the distance between the mouse’s head and the target center (Figure 1A). Thus, for electrode *i* with preferred direction *φ*_*i*_, stimulation frequency was calculated as:

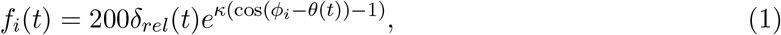

where *δ*_*rel*_(*t*) is the relative distance from the target at time *t* (scaling factor 0–1 to encode distance), *θ*(*t*) is the angle between subject’s forward heading and the vector from subject’s head to target, *κ* modulates the bandwidth of the curve, and the scalar 200 sets the maximum stimulation frequency at 200 Hz. For distal electrodes, 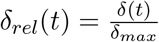 , while for proximal electrodes,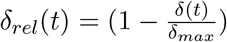.In all cases, *δ*_*max*_ = 25 cm (the diagonal distance across the behavioral training arena). The sparse encoding algorithm set *κ* = 16 and the redundant encoding algorithm set *κ* = 4.

**Analysis** *Logistic Regression*. We fit a log-likelihood generalized linear model to describe the probability of task success as a function of trial type (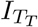 , baseline: ICMS), trial count (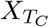 , baseline: 1), encoding algorithm (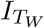 , baseline: sparse), training cohort (*I*_*C*_, baseline: cohort 1), sex of the animal (*I*_*S*_, baseline: female), and mouse ID (*I*_*M*_ , baseline: C1-1L1RF) with two-factor and three-factor interactions between trial count, encoding algorithm, and stimulation paradigm.

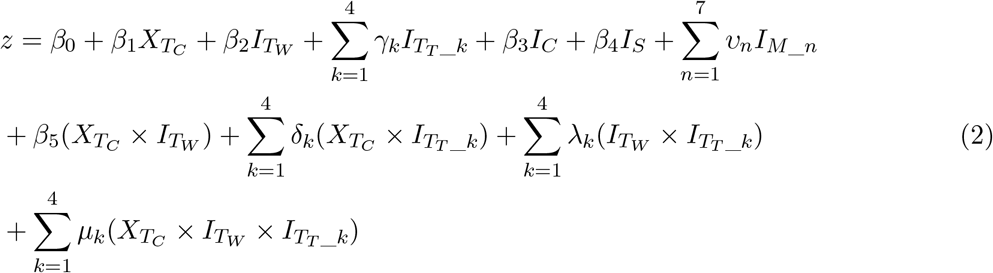

Probability of success was then calculated through the logit link function

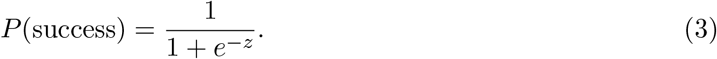

*Additional statistical testing* was performed using non-parametric statistics: the Wilcoxon signed-rank test for paired comparisons and the Wilcoxon rank-sum test for independent samples. Correction for multiple comparisons was made using the Holm-Bonferroni method.

*Quantification of single-trial behavior*. For all metrics excluding success rate, only successful trials were included in analyses.

*Success rate* was calculated as a window-moving average with a window size of 30 trials and 50% overlap between sequential windows. For analysis of the block switch and within-session switches, success rate was calculated as the fraction of trials in which the mouse successfully reached the target region within the allotted time in each session. For the within-session switches, we additionally included window-moving averages of success rate for multimodal, ICMS, and dim visual trials with a window of 4 trials and 50% overlap. We used a smaller window for this analysis because of the small proportion of trials performed for each tuning width during the within-session switches.

*Time-to-target* was measured as the total time between the start tone and the time the mouse’s head reaches the target. This provided a raw measure of how quickly mice find the target on average, regardless of starting distance from the target.

*Trial cropping*. On a subset of trials, the mouse was not facing the target at the start of the trial, impeding the immediate use of visual information. To fairly compare between visual and ICMS trials, we cropped behavioral trajectories on each trial to the point at which the mouse was first facing the target, defined as the point at which the mouse’s heading (vector between the nose and back of the head; Figure 1C) was within *±*123^*°*^ of the target direction, *δ*. Subsequent trial metrics were calculated using these cropped trajectories.

*Angular Dispersion* measures the spread of turning angles around an average direction of movement and provides a measure of path tortuosity: 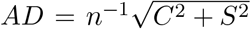 where 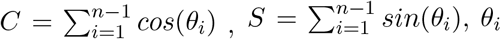 is the angle of movement between successive positions along the trajectory of movement and n is the number of data points collected along the path. A value of 0 denotes a circular path while a value of 1 denotes a straight path [59].

*Average Speed* was calculated as 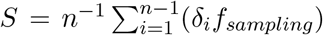 where *δ* is the distance between successive time points, *f*_*sampling*_ is the sampling rate (60 Hz), and n is the number of data points collected in a single trial.

## RESULTS

In this study, mice were trained to navigate to targets in a behavioral training cage, guided by sensory information that consisted of a visual target (dim visual and bright visual trials) and/or patterned multi-channel ICMS (ICMS trials, multimodal trials). We hypothesized that changing the encoding algorithm for the artificial sensory signal (redundant vs. sparse) would impact both learning rate and ICMS-guided performance. Across training sessions, visual trials provided an estimated timeline for task learning while ICMS trials provided an estimated timeline for learning of ICMS relative to task learning. Multimodal trials drive associative learning and later allow an assessment of multisensory integration of unimodal sensory information.

Mice were first split into two groups and assigned either sparse or redundant encoding (Figure 1D). This allowed us to compare learning speed and performance plateau across algorithms (Figure 2). Mice were considered to have “learned” the task when they reached proficiency, defined as a success rate of 75% on a single session; however, performance often fluctuated around this proficient level across sessions. Training on the initial ICMS algorithm continued until mice performed proficiently on five sequential sessions (an average of *≥*75% across days).

**Figure 2:**
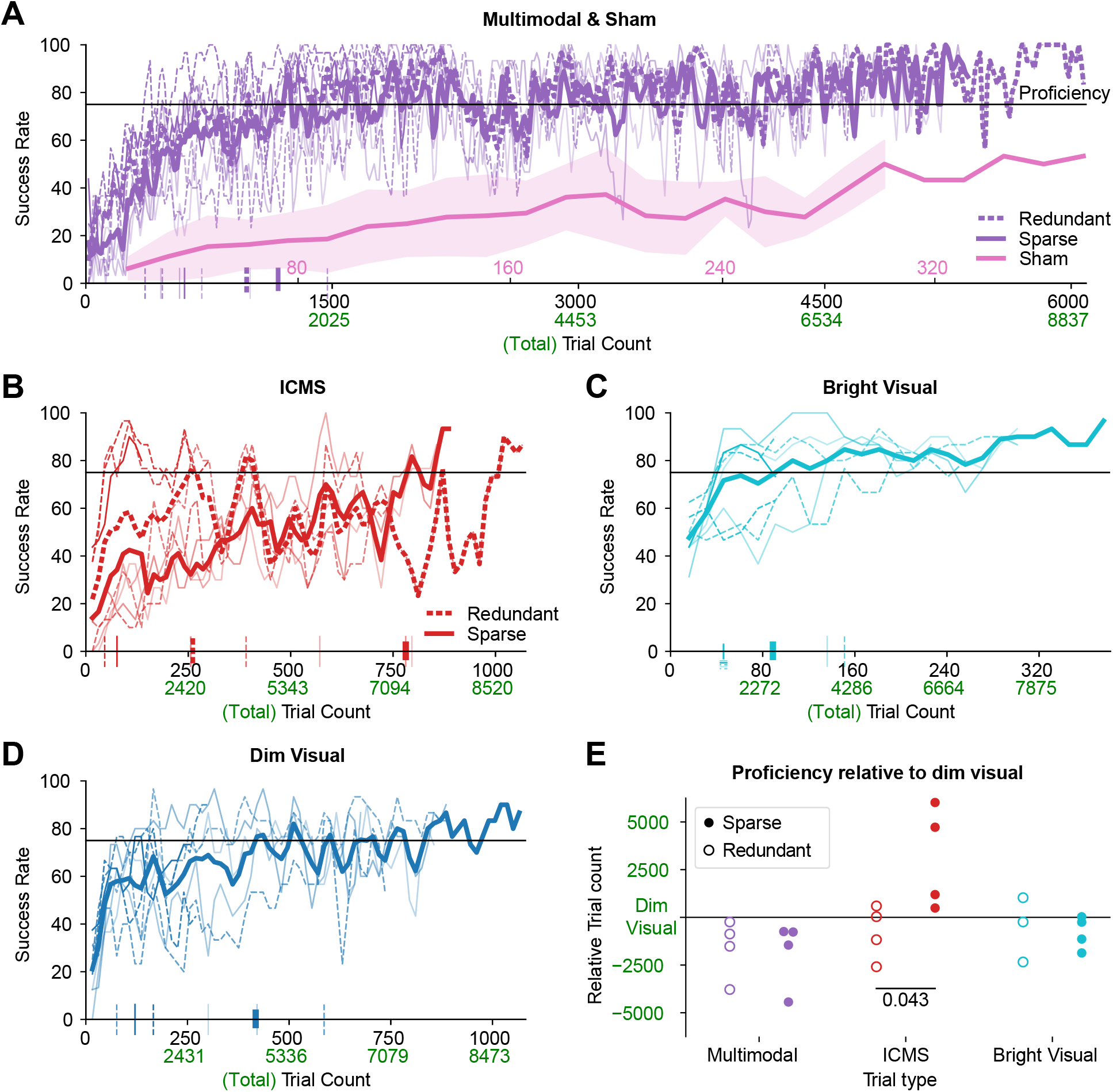
Moving average success rate on **A.** multimodal and sham, **B**. ICMS, **C**. bright visual, and **D**. dim visual trials. In all plots, solid lines denote sparse encoding while dashed denote redundant encoding with thin lines indicating individual mouse performance and thick lines indicating average performance. Separate sparse and redundant encoding averages are only shown for ICMS and multimodal trial types because these are the only trial types affected by the tuning width modulation. Time to reach proficiency, defined as the first 30 trial window to go above a 75% success rate, is indicated by ticks on the x-axis for each mouse. X-ticks in green indicate the total trial count (all trial types) at corresponding within-class trial count (black). **E**. Trial number at which proficiency is reached on each trial type relative to the trial at which proficiency was reached on dim visual trials. Mice trained with sparse encoding are shown filled circles and redundant encoding in open circles. Negative values indicate that proficiency was reached on that trial type prior to proficiency being reached on dim visual trials (e.g., multimodal and bright visual trials). Effect sizes between sparse and redundant are reported in Table **??** for each trial type. Comparison of ICMS proficiency between sparse and redundant encoding was performed with a Wilcoxon rank-sum test, as each mouse was first trained with a single encoding algorithm.

### Learning ICMS

We first consider performance with ICMS during the period of time mice were exposed to their initial encoding algorithm (Figure 2). We used a generalized linear model to relate task success to trial count, trial type, ICMS encoding algorithm, additionally including mouse sex, training cohort, and mouse ID as possible confounding variables (Eqn. 2). The model output is reported in full in Table **??**.

Many of the coefficients of the logistic regression model were significant predictors of success relative to baseline treatments (sparse ICMS learning in one female mouse in the first training cohort). First, all trial types (but sham) had higher success probability than ICMS trials. This shows that ICMS-guided trials had lower success rates across much of training than visually-guided or multimodal trials. Second, redundant encoding of ICMS shows performance gains relative to baseline sparse encoding. This model suggests our initial hypothesis was wrong; that, instead, redundant encoding improves ICMS-guided navigation relative to sparse encoding.

Next, to estimate a timeline for learning of the ICMS signal, we calculated a windowed moving-average of trial success rates on each trial type (Figure 2). Task proficiency on a particular trial type (multimodal, ICMS, etc.) was identified as the first window in which average success rates were *≥* 75%, shown for individual mice as vertical ticks along the x-axis in Figure 2. Mice rapidly learned the behavioral task, on average achieving proficiency on multimodal trials in fewer than 2000 total training trials (Figure 2A; total trials refers to the count of all trial types).

There was substantial variability in learning trajectories across mice, particularly between the two training cohorts, but also between male and female mice (Table **??**). To standardize the data across mice, we took the trial at which each mouse reached proficiency on dim visual trials as a reference. We then calculated the trial to reach proficiency for each mouse on the remaining trial types, and subtracted the dim visual reference point. This accounts for variability in task learning in individual mice and across training cohorts.

We found that mice took significantly longer to reach proficiency on ICMS trials with sparse than redundant encoding relative to when they reached proficiency on dim visual (p = 0.043, Wilcoxon-rank sum test; Figure 2E), suggesting that redundant encoding of sensory variables improves the learnability of ICMS-encoded artificial sensations. Further, mice reached proficiency on multimodal trials before dim visual trials (p= 0.015, paired t-test), which could be due to a superiority in performance with ICMS relative to vision or due to early multisensory integration that occurs even before the behavioral task is fully learned with natural vision. However, given that mice reach proficiency on multimodal trials at the same time regardless of ICMS encoding algorithm (p = 1, Wilcoxon rank-sum test), we conclude that the redundancy of the ICMS encoding algorithm has no effect on overall learning of the behavioral task.

### Block switch of ICMS encoding algorithms

Following the initial learning period (i.e., once mice became proficient using their original ICMS encoding algorithm), a mouse’s encoding algorithm was switched to the alternative code and kept fixed for the next five sessions (sparse encoding switched to redundant; redundant encoding switched to sparse, Figure 3). This allowed us to compare plateau performance on the two encoding algorithms for individual subjects. Note that plateau performance measures the animal’s ability to adapt to a particular encoding scheme on a longer timescale, such as hundreds of trials.

**Figure 3:**
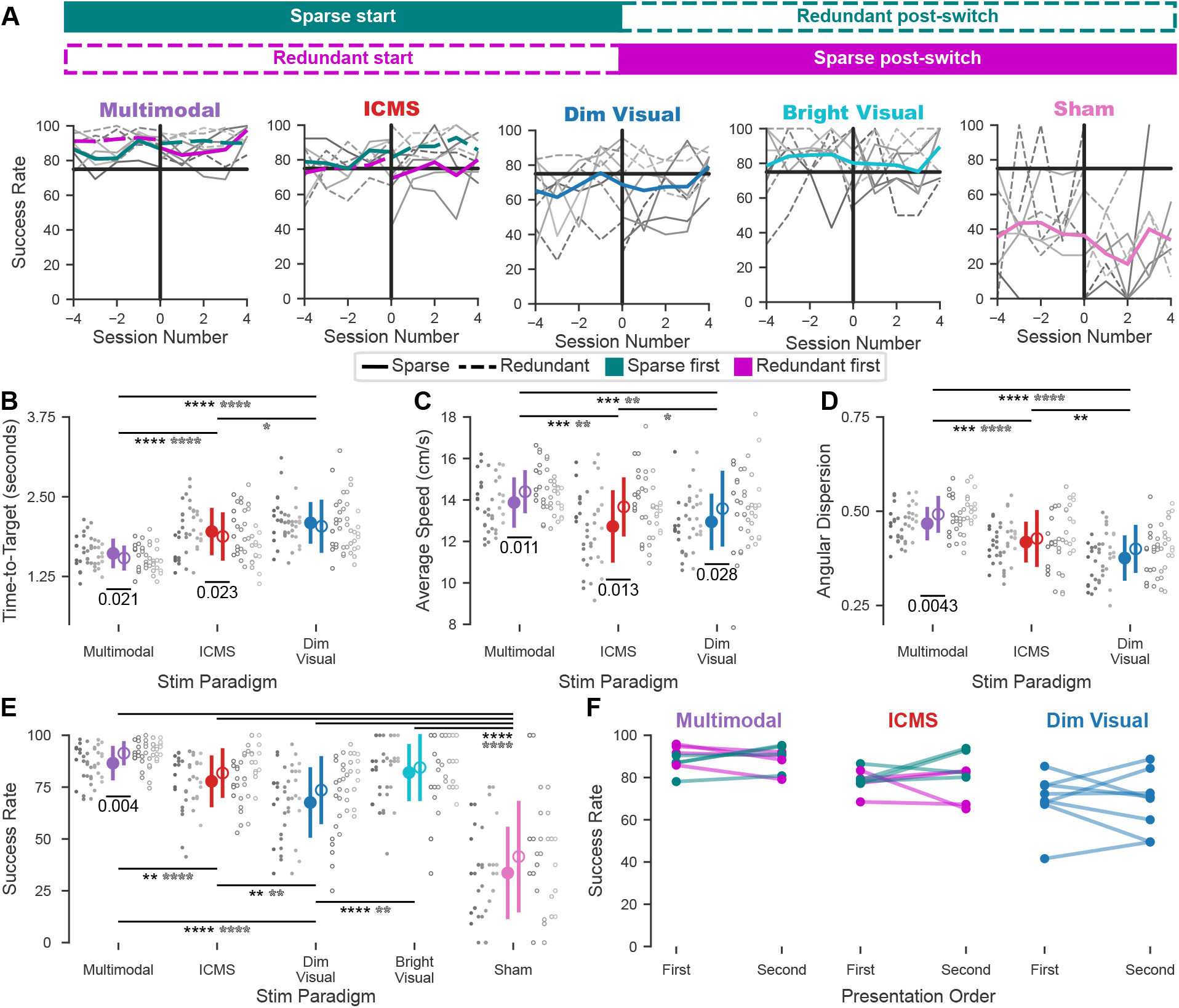
Task and ICMS learning. **A.** Average success rate (20-trial window; 50% overlap between windows) before and after the switch in encoding algorithms. Switch is at Session “0”. Line color indicates starting encoding algorithm (sparse start: teal, redundant start: magenta) and line style indicates current tuning width (sparse: solid, redundant: dashed). **B**. Average trial duration on successful trials (filled: sparse, open: redundant). Individual mouse performance is shown in gray with trial-type averages colored as in Figure 2. Significant differences across trial types for matched encoding algorithms are shown in filled/open stars for sparse/redundant comparisons. * *≤* p 0.05; ** p *≤* 0.01; *** p *≤* 0.001; **** p *≤* 0.0001 Colors, marker styles, and identification of significance are consistent across B-E. **C**. Average movement speed (cm/s) on successful trials. **D**. Average angular dispersion. **E**. Session-average success rates. **F**. Per-mouse success rates across five sessions pre- and post-switch. Individual mice are colored by starting encoding algorithm, consistent with **A**. There were no significant differences in success rate by presentation order on multimodal, ICMS, or dim visual trials.

A priori, any switch in ICMS encoding might be expected to impair ICMS-guided performance in trained mice. Although the encoding algorithms are similar in structure, the number of electrodes stimulating simultaneously differed substantially – two to three times as many during redundant encoding relative to sparse encoding. Therefore, mice switching from redundant to sparse encoding would suddenly experience a loss of information density, while mice switching from sparse to redundant encoding would suddenly experience a loss of specificity. Instead, as shown in Figure 3, mice showed little to no deficiency in performance following a change in encoding algorithm. There is a visible, but not significant, trend accompanying the switch (Figure 3): mice switching from redundant to sparse algorithms dropped an average of 4.6% in success rate on multimodal trials and 12.7% on ICMS trials. Mice switching from sparse to redundant increased an average of 3.03% in success rate on multimodal trials but decreased in ICMS performance by an average of 3.5%.

Despite a similar number of successful trials across encoding algorithms on ICMS-only trials (Figure 3A,E), we found significant differences in the quality of sensory-guided navigation across algorithms, as measured by time-to-target and average movement speed. Mice guided by redundant encoding made faster movements towards the target, resulting in shorter trial durations (Figure 3B,C). Further, redundant encoding facilitated integration of the ICMS signal with natural vision during multimodal trials. This was evident for the number of successful trials completed by mice (Figure 3E) but also appeared in movement quality, as evidenced by shorter trial durations, faster movement speed, and more directed movements towards the target during multimodal vs. uni-modal (ICMS or visual) trials (Figure 3B-D). These improvements could not be explained away by considering the order of the encoding algorithm mice experienced, as there was no net difference in performance between the first and second set of encoding algorithms (combining both algorithms; Figure 3F). Together, these results show that mice are able to decode the more redundant signal better than they are able to decode the sparse encoding signal, that redundant encoding facilitates multisensory integration of artificial sensory information with natural vision, and that neural circuits decoding artificial sensory information from ICMS are remarkably robust to changes in encoding redundancy.

### Within-session switching of ICMS encoding algorithms

To determine if there are intrinsic benefits to sparse vs. redundant encoding, outside of the context of across-session learning, we next implemented short, within-day switches of ICMS encoding (Figure 4). As these switches prohibit long-term adaptation, they better probe the facility by which neural circuits can directly interpret information encoded by patterned, multi-channel ICMS. This was achieved by training mice for five sessions in which the ICMS encoding algorithm was switched in the middle of the session (Figure 1D).

**Figure 4:**
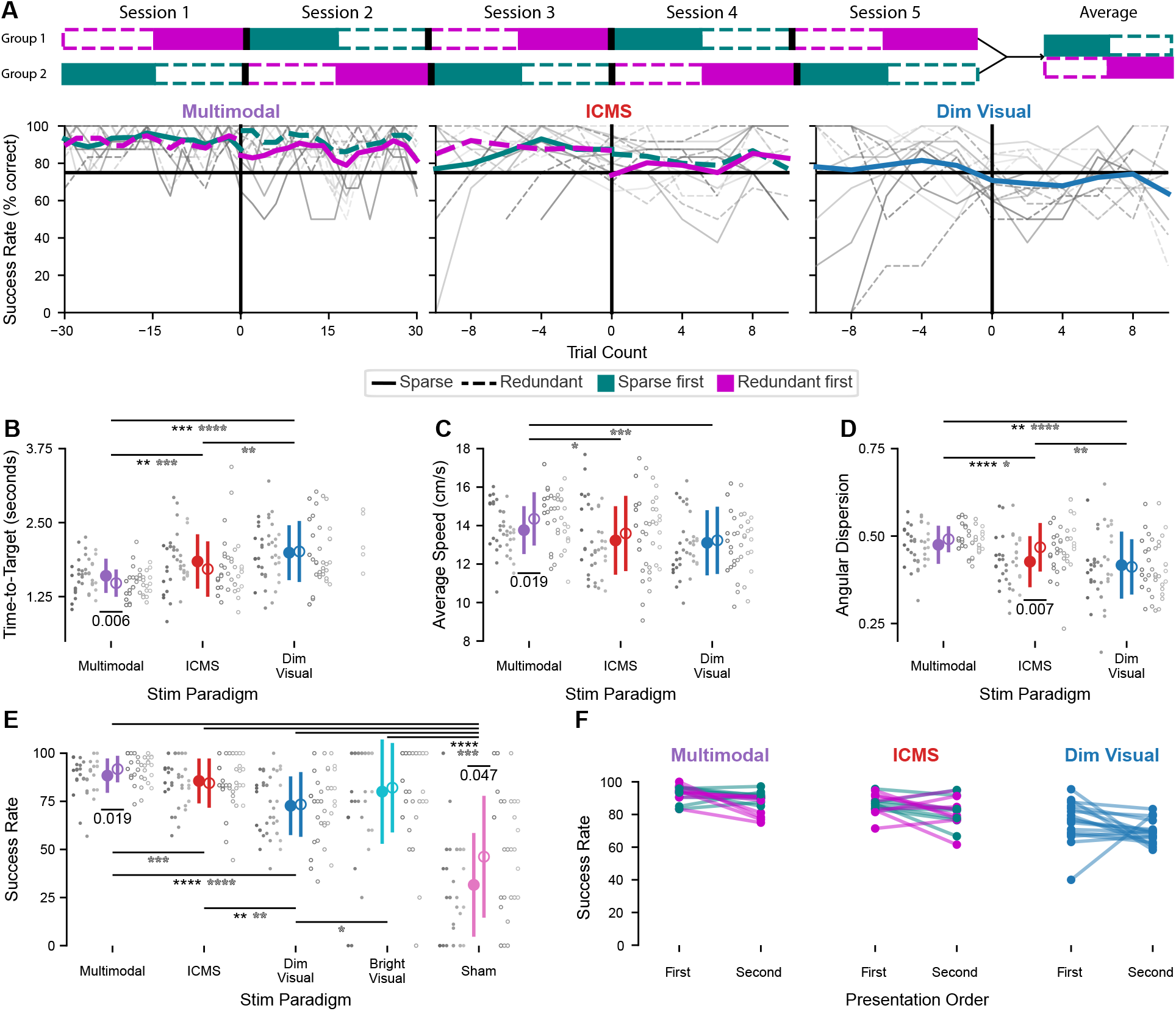
Within-session switches. **A.** Behavioral training structure for in-session switches (top). Below, four-trial rolling window average of success rates. Color indicates starting encoding algorithm (teal for sparse start and magenta for redundant start) and line style indicates encoding algorithm (solid for sparse, dashed for redundant). Gray success-rate lines show individual mouse performance averaged across sessions by starting encoding algorithm and colored lines show across-mouse averages by starting encoding algorithm (teal - sparse start, magenta - redundant start). **B**. Average trial duration on successful trials (filled: sparse, open: redundant). Individual mouse performance is shown in gray with trial-type averages colored as in Figure 2. Significant differences across trial types for matched encoding algorithms are shown in filled/open stars for sparse/redundant comparisons. * p *≤* 0.05; ** p *≤* 0.01; *** p *≤* 0.001; **** p *≤* 0.0001 Colors, marker styles, and identification of significance are consistent across B-E. **C**. Average speed on successful trials. **D**. Angular dispersion on successful trials. **E**. Average

We found that the outcome of within-session switches mirrored the results of block switches: mice did not struggle to switch between encoding algorithms, even at short timescales (Figure 4A). There did seem to be a minor effect of switching. On multimodal trials, success rate increased by 10%, on average, when switching from sparse to redundant encoding, and decreased by 7.9%, on average, when switching from redundant to sparse (Figure 4A,E). On ICMS trials, success rates dropped by *≥* 10% on average when switching from redundant to sparse. Still, as for block switches, the effects were not significant.

Next, we averaged performance pre- and post-switch to compare across trial types and encoding algorithms (Figure 4B-E). On unimodal ICMS trials, the only difference between sparse and redundant encoding was in angular dispersion, with more direct movements with redundant encoding (Figure 4D). Stable performance within a session suggests that mice were able to retain and flexibly switch between stimulation protocols. Once again, significant differences were apparent on multimodal trials. Performance on multimodal trials was superior with redundant encoding relative to sparse encoding in terms of success rate, time-to-target, and average movement speed (Figure 4B,C,E). However, in contrast to block switches, integration was weaker for sparse ICMS encoding during within-session switches, as there was no significant difference in performance between sparse multimodal and sparse ICMS trials on half of the performance measures (Figure 4B-E). This supports our prior results showing that redundant encoding of ICMS facilitates integration of natural and artificial sensory information.

Differences between sparse and redundant within-session periods were not caused by the order in which mice were presented with each stimulating algorithm, as there were no significant differences in performance when simply considering the first and second stimulation protocol, despite a weak trend toward poorer performance in the second half of each training session (Figure 4F). To address the question of whether there are changes in motivation across the training session that impact performance, we additionally analyzed success of dim visual trials during the first and second half of each session (Figure 4F). As for ICMS and multimodal trials, there was a weak decrease in dim visual performance over the course of a session, but the change was insignificant. Therefore, motivation remains relatively stable across the first and second half of the behavioral training sessions. success rate by mouse, trial type, and encoding algorithm. **F**. Average success rate for each mouse as a function of starting encoding algorithm (indicated by marker/line color, as in A) and presentation order for multimodal, ICMS, and dim visual trials. Colors are consistent with A.

## DISCUSSION

### Learning artificial sensations

The optimal approach to encoding artificial sensation is still uncertain. There are clear limits to how precisely we can use electrical microstimulation to activate neurons [19, 60–64], and in turn, how those patterns of neural activation elicit certain percepts [11, 21, 32, 34]. However, there are likely also limits to a patient’s ability to learn consistent patterns in ICMS as sensory feedback in the context of learning-based artificial sensation [35, 65, 66]. Our current efforts very much fall on the side of leveraging learning to encode artificial sensory feedback. We showed here that, not only can freely-moving mice rapidly learn and integrate an ICMS signal with natural vision [43], but also: 1) the stimulation algorithm can be fine-tuned to improve both learning speed and plateau performance, 2) the improvement of redundant encoding over sparse encoding is evident at both short and long timescales, and 3) decoding of learned ICMS algorithms is surprisingly flexible and robust, with only minor impairment even just after a switch.

### Sparse versus redundant encoding algorithms

The engineering question of interest was how to increase the information encoded by patterned, multi-channel ICMS of the cortex and to promote the integration of artificial sensation into existing neural circuits processing sensorimotor function. We addressed this here by manipulating the algorithm used to encode artificial sensation — providing mice either sparse or redundant encoding, and testing which was easiest to learn and integrate with natural vision. There are trade-offs to both algorithms. The benefit of sparse encoding is that activations are more precise and spatially localized on the brain, and that a fixed change in the encoded variable (direction, *θ*) leads to a larger change in stimulation frequency than for the redundant code [52]. Such specificity could be easier to learn and interpret [47], but it comes at the cost of naturally being a weaker signal, given that a smaller population of neurons are activated. In contrast, redundant encoding activates a larger total population of neurons, yielding a stronger signal that is easier to detect [67, 68], but comes at the cost of spatial specificity, which impairs the discriminability of stimulation between two distinct electrode sites and yields smaller changes in stimulation frequency for a fixed change in *θ* (Figure **??**). Increasing the amount of stimulation the mouse receives could also have over-excited the neural population, impairing behavior and even causing seizures [69]. We note that structural similarity between the two encoding algorithms, where one is a stretched version of the other, also probably facilitated switching between algorithms. There are other ways to implement sparsity and redundancy, and the outcome of these experiments may, in part, have to do with the details of our implementation.

### Encoding algorithms for artificial sensation: what does it mean to be biomimetic?

Stimulation algorithms that are considered biomimetic are those that identify and selectively stimulate subpopulations of neurons to replicate the neural activity (and, theoretically, the perceptual quality) associated with natural sensations [11, 12, 20, 21, 33, 34]. While this approach does, in fact, elicit naturalistic sensations, they are limited by the features of neurons proximal to the electrode contact sites. In general, any electrode array with an arbitrary, fixed number of electrode sites will have limited access to the full set of sensory information that is required to provide real-time information describing limb movements [70], suggesting that some learning or adaptation will be required to precisely map artificial sensory feedback to artificial motor control. Therefore, our current explorations of the limits of neural plasticity (learning) in the context of encoding artificial sensation [71] can be viewed as providing a lower bound for artificial information content.

Learning-based stimulation is superficially the opposite of biomimetic stimulation; however, the encoding algorithms we used directly mimic neural activity during natural sensory processing. In fact, we treat electrodes as if they are sensory neurons encoding stimuli, assigning them stimulation algorithms that mimic response properties of sensory neurons. For example, neurons in primary somatosensory cortex of the mouse show classical bell-shaped tuning to passive movement direction and neural responses scale linearly with limb movement distance [55]. We employ precisely these relationships in the ICMS encoding algorithms described in Figure 1. This code imposes a fixed relationship between sensor input (animal heading) and ICMS output (stimulation across 16 electrodes) that subjects can then learn through experience performing movements. Multiple animal models have been capable of learning such fixed mappings [25, 26, 28, 29, 45, 46, 72–74]. Further, behavior guided by patterned ICMS not only rivals that with natural vision [43, 72] but is further improved under multisensory conditions. Thus, not only can learned artificial sensation be as reliable as natural sensation, the integration of the new signal into neural circuits governing sensorimotor function is rapid and complete.

One interpretation of our results aligns with a modern view that the brain is continuously predicting incoming sensory consequences of the body’s actions [75–78]. Like ICMS in this study, prediction of natural sensory consequences of movement must be learned, as the neural circuits required must be developed through experience [79]. When the structure of the sensorimotor control problem changes, such as during tool or prosthetics use (both of which provide a novel input/output relationship), neural circuits adapt accordingly [35, 38, 39, 80]. To help neural circuits predict and adapt, an encoding algorithm should impose a fixed relationship between sensory variables and stimulation parameters. This will allow animals and humans to not only learn an association between artificial sensation and motor output, but even to predict upcoming artificial percepts.

## References

[1] Brian S. Armour, Elizabeth A. Courtney-Long, Michael H. Fox, Heidi Fredine, and Anthony Cahill. “Prevalence and causes of paralysis - United States, 2013”. In: American Journal of Public Health 106.10 (2016), pp. 1855–1857.

[2] 111th U.S. Congress. Christopher and Dana Reeve Paralysis Act. House Report No. 111–45. 2009.

[3] Henri Lorach et al. “Walking naturally after spinal cord injury using a brain–spine interface”. In: Nature 618.7963 (2023), pp. 126–133.

[4] Claudia A. Angeli, V. Reggie Edgerton, Yury P. Gerasimenko, and Susan J. Harkema. “Altering spinal cord excitability enables voluntary movements after chronic complete paralysis in humans”. In: Brain 137.5 (2014), pp. 1394–1409.

[5] R van den Brand et al. “Restoring voluntary control of locomotion after paralyzing spinal cord injury”. In: Science 336.6085 (2012), pp. 1182–1185.

[6] Jose M. Carmena et al. “Learning to control a brain-machine interface for reaching and grasping by primates”. In: PLoS Biology 1.2 (2003).

[7] Amy L. Orsborn et al. “Closed-loop decoder adaptation shapes neural plasticity for skillful neuroprosthetic control”. In: Neuron 82.6 (2014), pp. 1380–1393.

[8] Krishna V. Shenoy and Jose M. Carmena. “Combining decoder design and neural adaptation in brain-machine interfaces”. In: Neuron 84.4 (2014), pp. 665–680.

[9] D. Young et al. “Closed-loop cortical control of virtual reach and posture using Cartesian and joint velocity commands”. In: Journal of Neural Engineering 16.2 (2019).

[10] Sharlene N. Flesher et al. “A brain-computer interface that evokes tactile sensations improves robotic arm control”. In: Science 372.May (2021), pp. 831–836.

[11] Giacomo Valle et al. “Tactile Edges and Motion Via Patterned Microstimulation of the Human Cortex”. In: Science 387 (2025), pp. 315–322.

[12] Michelle Armenta Salas et al. “Proprioceptive and cutaneous sensations in humans elicited by intracortical microstimulation”. In: eLife 7 (2018), pp. 1–11.

[13] A. J. Suminski, D. C. Tkach, A. H. Fagg, and N. G. Hatsopoulos. “Incorporating Feedback from Multiple Sensory Modalities Enhances Brain-Machine Interface Control”. In: Journal of Neuroscience 30.50 (2010), pp. 16777–16787.

[14] Philip N. Sabes. “Sensory integration for reaching. Models of optimality in the context of behavior and the underlying neural circuits”. In: Progress in Brain Research 191 (2011), pp. 195–209.

[15] Marc O Ernst and Martin S Banks. “Humans integrate visual and haptic information in a statistically optimal fashion.” In: Nature 415.6870 (2002), pp. 429–433. eprint: NIHMS150003.

[16] Barry E Stein and Terrence R Stanford. “Multisensory integration: current issues from the perspective of the single neuron.” In: Nature reviews. Neuroscience 9.4 (2008), pp. 255–266.

[17] Isabelle A. Rosenthal et al. “Visual context affects the perceived timing of tactile sensations elicited through intra-cortical microstimulation”. In: bioRxiv (2024). eprint: https://www.biorxiv.org/content/early/2024/05/14/2024.05.13.593529.full.pdf.

[18] Sergejus Butovas and Cornelius Schwarz. “Spatiotemporal Effects of Microstimulation in Rat Neocortex: A Parametric Study Using Multielectrode Recordings”. In: J Neurophysiol. 90 (2003), pp. 3024–3039.

[19] Mark H Histed, Vincent Bonin, and R Clay Reid. “Direct activation of sparse, distributed populations of cortical neurons by electrical microstimulation.” In: Neuron 63.4 (2009), pp. 508–22.

[20] Gregg a Tabot et al. “Restoring the sense of touch with a prosthetic hand through a brain interface.” In: Proc. Natl. Acad. Sci. USA 110.45 (2013), pp. 18279–84.

[21] S. N. Flesher et al. “Intracortical microstimulation of human somatosensory cortex”. In: Sci Transl Med. 8.361 (2016), 361ra141–361ra141.

[22] Thierri Callier, Nathan W Brantly, Attilio Caravelli, and Sliman J Bensmaia. “The frequency of cortical microstimulation shapes artificial touch”. In: Proc Natl Acad Sci USA 117.2 (2020), pp. 1191–1200.

[23] Christopher Hughes and Takashi Kozai. “Dynamic amplitude modulation of microstimulation evokes biomimetic onset and offset transients and reduces depression of evoked calcium responses in sensory cortices”. In: Brain Stimulation 16 (2023), pp. 939–965.

[24] Maria C. Dadarlat, Joseph E. O’Doherty, and Philip N. Sabes. “A learning-based approach to artificial sensory feedback leads to optimal integration”. In: Nat Neurosci. 18.1 (2015), pp. 138–144.

[25] Maria C. Dadarlat and Philip N. Sabes. “Encoding and Decoding of Multi-Channel ICMS in Macaque Somatosensory Cortex”. In: IEEE Trans Haptics 9.4 (2016), pp. 508–514.

[26] Eric E. Thomson et al. “Cortical neuroprosthesis merges visible and invisible light without impairing native sensory function”. In: eNeuro 4.6 (2017), pp. 1–17.

[27] Konstantin Hartmann et al. “Embedding a panoramic representation of infrared light in the adult rat somatosensory cortex through a sensory neuroprosthesis”. In: Journal of Neuroscience 36.8 (2016), pp. 2406–2424.

[28] Sanjiv K. Talwar et al. “Rat navigation guided by remote control”. In: Nature 417.6884 (2002), pp. 37–38.

[29] Andrew G. Richardson et al. “Learning active sensing strategies using a sensory brain–machine interface”. In: Proceedings of the National Academy of Sciences 116.35 (2019), p. 201909953.

[30] John S Choi et al. “Eliciting naturalistic cortical responses with a sensory prosthesis via optimized microstimulation”. In: Journal of Neural Engineering 13.5 (2016), p. 056007.

[31] Karthik Kumaravelu et al. “A comprehensive model-based framework for optimal design of biomimetic patterns of electrical stimulation for prosthetic sensation”. In: J Neural Eng. 17.4 (2020), p. 046045.

[32] Christopher L. Hughes et al. “Perception of microstimulation frequency in human somatosensory cortex”. In: eLife 10 (2021), pp. 1–19.

[33] Charles M. Greenspon, Natalya D. Shelchkova, Taylor G. Hobbs, Sliman J. Bensmaia, and Robert A. Gaunt. “Intracortical microstimulation of human somatosensory cortex induces natural perceptual biases”. In: Brain Stimulation 17.6 (2024), pp. 1178–1185.

[34] Taylor G. Hobbs et al. “Biomimetic stimulation patterns drive natural artificial touch percepts using intracortical microstimulation in humans”. In: Journal of Neural Engineering 22.3 (2025).

[35] Ivana Cuberovic, Anisha Gill, Linda J. Resnik, Dustin J. Tyler, and Emily L. Graczyk. “Learning of Artificial Sensation Through Long-Term Home Use of a Sensory-Enabled Prosthesis”. In: Frontiers in Neuroscience 13.August (2019), pp. 1–24.

[36] Robin Kim and Lan Luan. Ultraflexible Electrodes for Low-Threshold, High-Resolution Microstimulation: Chronic Stability and Neuronal Excitability Changes. Chicago, IL: Society for Neuroscience, 2024.

[37] G. H. Recanzone, M. M. Merzenich, W. M. Jenkins, K. A. Grajski, and H. R. Dinse. “Topographic reorganization of the hand representation in cortical area 3b of owl monkeys trained in a frequency-discrimination task”. In: Journal of Neurophysiology 67.5 (1992), pp. 1031–1056.

[38] Solaiman Shokur et al. “Expanding the primate body schema in sensorimotor cortex by virtual touches of an avatar”. In: Proceedings of the National Academy of Sciences of the United States of America 110.37 (2013), pp. 15121–15126.

[39] Angelo Maravita and Atsushi Iriki. “Tools for the body (schema)”. In: Trends in Cognitive Sciences 8.2 (2004), pp. 79–86.

[40] Joseph G. Makin, Matthew R. Fellows, and Philip N. Sabes. “Learning Multisensory Integration and Coordinate Transformation via Density Estimation”. In: PLoS Computational Biology 9.4 (2013), e1003035.

[41] Liping Yu, Benjamin a Rowland, and Barry E Stein. “Initiating the development of multisensory integration by manipulating sensory experience.” In: The Journal of neuroscience : the official journal of the Society for Neuroscience 30.14 (2010), pp. 4904–4913.

[42] Barry E Stein, Terrence R Stanford, and Benjamin A Rowland. “Development of multisensory integration from the perspective of the individual neuron”. In: Nature Reviews Neuroscience 15.8 (2014), pp. 520–535.

[43] Samuel Senneka and Maria C. Dadarlat. “Integration of learned artificial sensation with vision during freely-moving navigation”. In: Proceedings of the National Academy of Sciences of the United States of America (9 2026), e2521769123.

[44] Eric E. Thomson, Rafael Carra, and Miguel A.L. Nicolelis. “Perceiving invisible light through a somatosensory cortical prosthesis”. In: Nature Communications 4 (2013), p. 1482. eprint: NIHMS150003.

[45] Henri Lassagne et al. “Continuity within the somatosensory cortical map facilitates learning”. In: Cell Reports 39.1 (2022), p. 110617.

[46] Aamir Abbasi et al. “Brain-machine interface learning is facilitated by specific patterning of distributed cortical feedback”. In: Science Advances 9.38 (2023).

[47] Thomas J. Smith et al. “Investigating the spatial limits of somatotopic and depth-dependent sensory discrimination stimuli in rats via intracortical microstimulation”. In: Frontiers in Neuroscience 19 (May 2025), p. 1602996.

[48] Morgan E. Urdaneta, Nicolas G. Kunigk, Francisco Delgado, Shelley I. Fried, and Kevin J. Otto. “Layer-specific parameters of intracortical microstimulation of the somatosensory cortex”. In: Journal of Neural Engineering 18.5 (2021).

[49] Andrew S Koivuniemi, Kevin J Otto, and Abstract Intracortical. “Asymmetric versus symmetric pulses for cortical microstimulation”. In: IEEE TRANSACTIONS ON NEURAL SYSTEMS AND REHABILITATION ENGINEERING 19.5 (2011), pp. 468–476.

[50] Bruno A. Olshausen and David J. Field. “Sparse coding of sensory inputs”. In: Current Opinion in Neurobiology 14.4 (2004), pp. 481–487.

[51] Matthew Chalk and Olivier Marre. “Toward a unified theory of efficient , predictive , and sparse coding”. In: PNAS 115.1 (2018), pp. 186–191.

[52] Alexandre Pouget and Peter E Latham. “Narrow Versus Wide Tuning Curves : What ‘s Best for a Population Code ?” In: Neural Computation 11 (1999), pp. 85–90.

[53] Yu Cheng Pei, Steven S. Hsiao, James C. Craig, and Sliman J. Bensmaia. “Shape invariant coding of motion direction in somatosensory cortex”. In: PLoS Biology 8.2 (Feb. 2010), e1000305.

[54] M J Prud’homme and J F Kalaska. “Proprioceptive activity in primate primary somatosensory cortex during active arm reaching movements.” In: Journal of neurophysiology 72.5 (1994), pp. 2280–301.

[55] Ignacio Alonso et al. “Peripersonal encoding of forelimb proprioception in the mouse somatosensory cortex”. In: Nature Communications 14 (2023), p. 1866.

[56] Authors David A Bjånes, Luke Bashford, Kelsie Pejsa, Brian Lee, and Y Charles. “Multi-channel intra-cortical micro-stimulation yields quick reaction times and evokes natural somatosensations in a human participant”. In: medRxiv (2022).

[57] George Paxinos and Keith B. J. Franklin. Paxinos and Franklin’s the Mouse Brain in Stereotaxic Coordinates. 5th. Academic Press, 2019.

[58] Gary A Kane, Jonny L Saunders, and Alexander Mathis. “Real-time, low-latency closed-loop feedback using markerless posture tracking”. In: eLife 9 (2021), e61909.

[59] Curtis Miller, Mary C. Christman, and Inma Estevez. “Movement in a confined space: Estimating path tortuosity”. In: Applied Animal Behaviour Science 135 (1-2 Nov. 2011), pp. 13–23.

[60] Karthik Kumaravelu and Warren M. Grill. “Neural mechanisms of the temporal response of cortical neurons to intracortical microstimulation”. In: Brain Stimulation 17.2 (2024), pp. 365–381.

[61] Karthik Kumaravelu, Joseph Sombeck, Lee E. Miller, Sliman J. Bensmaia, and Warren M. Grill. “Stoney vs. Histed: Quantifying the spatial effects of intracortical microstimulation”. In: Brain Stimulation 15.1 (2022), pp. 141–151.

[62] Maria C. Dadarlat, Yujiao Jennifer Sun, and Michael P. Stryker. “Activity-dependent recruitment of inhibition and excitation in the awake mammalian cortex during electrical stimulation”. In: Neuron 112.5 (2024), 821–834.e4.

[63] Christopher L Hughes, Kevin C Stieger, Keying Chen, Alberto L Vazquez, and Takashi D Y Kozai. “Spatiotemporal properties of cortical excitatory and inhibitory neuron activation by sustained and bursting electrical microstimulation”. In: iScience 28.6 (2025), p. 112707.

[64] Richy Yun, Jonathan H. Mishler, Steve I. Perlmutter, Rajesh P. N. Rao, and Eberhard E. Fetz. “Responses of cortical neurons to intracortical microstimulation in awake primates”. In: eNeuro 10.4 (2023), ENEURO.0336–22.2023.

[65] Silvia L Isabella et al. “Artificial embodiment displaces cortical neuromagnetic somatosensory responses”. In: Scientific Reports 14 (2024), p. 22279.

[66] Daan B. Wesselink et al. “Obtaining and maintaining cortical hand representation as evidenced from acquired and congenital handlessness”. In: eLife 8 (2019), pp. 1–19.

[67] Boubker Zaaimi, Ricardo Ruiz-Torres, Sara a Solla, and Lee E Miller. “Multi-electrode stimulation in somatosensory cortex increases probability of detection.” In: Journal of Neural Engineering 10.5 (Oct. 2013), p. 056013. eprint: NIHMS150003.

[68] Nicolas G. Kunigk, Morgan E. Urdaneta, Ian G. Malone, Francisco Delgado, and Kevin J. Otto. “Reducing Behavioral Detection Thresholds per Electrode via Synchronous, Spatially-Dependent Intracortical Microstimulation”. In: Frontiers in Neuroscience 16.June (2022), pp. 1–11.

[69] Naofumi Suematsu, Alberto L Vazquez, and Takashi Dy. “Chronic alteration of Ca 2 + and hemodynamic signals induced by intracortical microstimulation in the visual cortex of awake mice”. In: Biomaterials 334.May (2026), p. 124276.

[70] Robin Kim, Yuxuan Liu, Jiaao Zhang, and Chong Xie. “Towards precise synthetic neural codes : high-dimensional stimulation with fl exible electrodes”. In: npj Flexible Electronics 9.68 (2025).

[71] Maria C Dadarlat, Ryan A Canfield, and Amy L Orsborn. “Neural Plasticity in Sensorimotor Brain-Machine Interfaces”. In: Annual Review of Biomedical Engineering 15.2 (2023), pp. 51–76.

[72] Maria C Dadarlat. “Artificial Sensory Feedback for Neural Prostheses”. PhD thesis. University of California, San Francisco, 2014, pp. 1078–1085.

[73] Joseph E. O’Doherty, Solaiman Shokur, Leonel E. Medina, Mikhail A. Lebedev, and Miguel A.L. Nicolelis. “Creating a neuroprosthesis for active tactile exploration of textures”. In: Proceedings of the National Academy of Sciences of the United States of America 116.43 (2019), pp. 21821–21827.

[74] Aamir Abbasi, Dorian Goueytes, Daniel E. Shulz, Valérie Ego-Stengel, and Luc Estebanez. “A fast intracortical brain-machine interface with patterned optogenetic feedback”. In: Journal of Neural Engineering 15.4 (2018).

[75] Laurence Aitchison and Máté Lengyel. “With or without you: predictive coding and Bayesian inference in the brain”. In: Current Opinion in Neurobiology 46 (2017), pp. 219–227.

[76] Rajesh P.N. Rao and Dana H. Ballard. “Predictive coding in the visual cortex: A functional interpretation of some extra-classical receptive-field effects”. In: Nature Neuroscience 2.1 (1999), pp. 79–87.

[77] Georg B. Keller and Thomas D. Mrsic-Flogel. “Predictive Processing: A Canonical Cortical Computation”. In: Neuron 100.2 (2018), pp. 424–435.

[78] Rajesh P.N. Rao. “A sensory–motor theory of the neocortex”. In: Nature Neuroscience 27.June 2023 (2024).

[79] Alexander Attinger, Bo Wang, and Georg B. Keller. “Visuomotor Coupling Shapes the Functional Development of Mouse Visual Cortex”. In: Cell 169.7 (2017), 1291–1302.e14.

[80] Zineb Hayatou, Hongkai Wang, Antoine Chaillet, Daniel E Shulz, and Valérie Ego-stengel. “Embodiment of an artificial limb in mice”. In: PLOS Biology 23.6 (2025), e3003186.

